# Effects of learning and escitalopram administration on serum BDNF levels, a randomised placebo-controlled trial

**DOI:** 10.1101/2021.04.09.439014

**Authors:** J Unterholzner, GM Godbersen, M Klöbl, V Ritter, D Pacher, R Seiger, N Varghese, A Eckert, R Lanzenberger, T Vanicek

## Abstract

**Background:** The brain-derived neurotrophic factor (BDNF) has been implicated in processes essential for neuroplasticity. Learning and serotonin reuptake inhibitors (SSRI) foster neuronal reorganization, a mechanism potentially related to BDNF. This study aims to assess the effects of associative learning and escitalopram on serum BDNF (sBDNF) levels, to gain further knowledge on their dynamic interplay.

**Methods:** For three weeks, 37 participants performed one of two associative learning paradigms with either emotional or semantic content daily. During a subsequent three-week period of reversal learning, subjects either received escitalopram (10mg per day) or placebo. Before and after each learning period sBDNF values were assessed. Citalopram plasma levels were measured at the last time point. Linear mixed effects models (LME) and partial Spearman’s rank and Pearson correlations were used for statistical analyses.

**Results:** One-way LME resulted in a significant effect of time during the first learning period over both groups (p<0.01). Two-way LME revealed a significant interaction effect of the emotional content learning group and time (p=0.02). Three-way LME (time x reversal learning group x substance) showed no significant effects (all p> 0.05). Furthermore, correlation between citalopram and sBDNF level after three weeks of escitalopram administration exhibit a negative trend (partial Pearson correlation: r=-0.30, p=0.05; partial Spearman’s rank: r=-0.22, p=0.15).

**Conclusion:** The results suggest that three weeks of associative emotional content learning affect sBDNF levels, while subsequently assessed citalopram plasma and sBDNF levels tend to correlate negatively.

**Key Points:**

- Emotional learning may affect serum BDNF levels in healthy human subjects
- Blood levels of citalopram and serum BDNF exhibit a negative correlation

## 1. Introduction

The brain-derived neurotrophic factor (BDNF) is a protein of the neurotrophin family that has been implicated in neuroplastic processes and linked with brain disorders[1]. Due to its effects on neuronal differentiation, axon-dendrite growth and guidance, synapse formation and the modulation of neuronal activation[2], BDNF could also be proposed as a surrogate marker for neuroplasticity. The neurotrophin is highly expressed in the central nervous system (CNS), particularly in the hippocampus, but it can be determined in blood, as well. While the exact origin of circulating BDNF outside of the central nervous system is not yet fully understood[3], current evidence suggests that BDNF can be transported through the blood-brain barrier[4]. BDNF concentrations in the brain, particularly in the prefrontal cortex and hippocampus[5], are linked with BDNF serum levels. Also, peripherally administered BDNF elicits changes in the CNS, suggesting that it might be relevant for brain metabolism and function.

Through its neuroplastic properties, BDNF has repeatedly been associated with learning. For example, BDNF expression positively influences spatial memory in mice[6, 7] and contributes to long-term potentiation (LTP), a process that is critical for memory consolidation and learning[2]. Moreover, levels of BDNF appear to decline with hippocampal volume and memory[8] and BDNF seems to be especially relevant for hippocampal-dependent learning[9].

Increasing evidence suggests that BDNF and neuroplastic mechanisms can also be influenced through selective serotonin reuptake inhibitors (SSRIs)[10]. Chronic SSRI administration activates cAMP-response element binding protein (CREB), a transcription factor, which in turn favours synaptic restructuring. Since the BDNF gene has a cAMP-response element, SSRIs might thus influence BDNF expression through activation of CREB[11]. BDNF also unfolds its effect via the tropomyosin-related kinase B (TrkB) receptor and downstream signalling cascades[2]. Recently, the group around Eero Castrén reported that the SSRI fluoxetine increased the accessibility of TrkB for BDNF by directly binding to the TrkB transmembrane domain[12]. The authors remark that clinical response to an antidepressant requires high enough brain concentrations to bind TrkB, which is understood to present a low-affinity target for serotonergic agents. SSRIs influence mood[13] and motor function after ischemic stroke[14] and affect fear extinction learning[15] as well as contingency learning[16], all of which might be ascribed to the modulation of neuroplastic processes involving BDNF and TrkB. Interestingly, administration of BDNF alone also influences fear extinction[17]. If, in turn, BDNF can be influenced by learning alone has yet to be investigated in healthy human subjects. Moreover, given the molecular foundation, one might expect a change of central, but also peripheral BDNF levels through chronic SSRI administration. However, data on this topic are still scarce.

To unravel this dynamic interplay between learning, chronic SSRI administration and peripheral BDNF levels in healthy humans, we performed an intervention-study using an associative learning task and a subsequent associative relearning period with concomitant daily administration of either placebo or SSRI. We aimed to investigate, i) if sBDNF levels change dependent on different learning groups (emotional/semantic content) over time, ii) if there is an effect of SSRI administration and associative reversal learning on sBDNF levels over time and iii) if there is a correlation between sBDNF levels and citalopram plasma levels.

## 2. Material and Methods

### Study design

This present analysis is part of a larger randomized placebo-controlled, double-blinded intervention study in healthy human subjects, approved by the ethics committee of the Medical University of Vienna (EK Nr.: 1739/2016). It has been performed in accordance with the Declaration of Helsinki (1964). The project is registered at clinicaltrials.gov with the identifier NCT02753738.

After recruitment and inclusion, subjects were randomly assigned to one of two groups for associative learning, either with emotional (face pairs) or semantic content (Chinese characters and unrelated German nouns). At three time points, serum levels of BDNF were assessed: at baseline (m1), after a period of associative learning (m2) and again after a period of associative reversal learning (m3). Before the associative reversal learning period, subjects were randomly assigned to one of two substance groups, either 10mg of escitalopram (SSRI) daily, or a daily intake of placebo drug (see Fig 1). Both, participants and research staff were blinded for substance group. Additionally, citalopram plasma levels were assessed during the three-week study drug administration and at time point m3.

**Figure 1.**
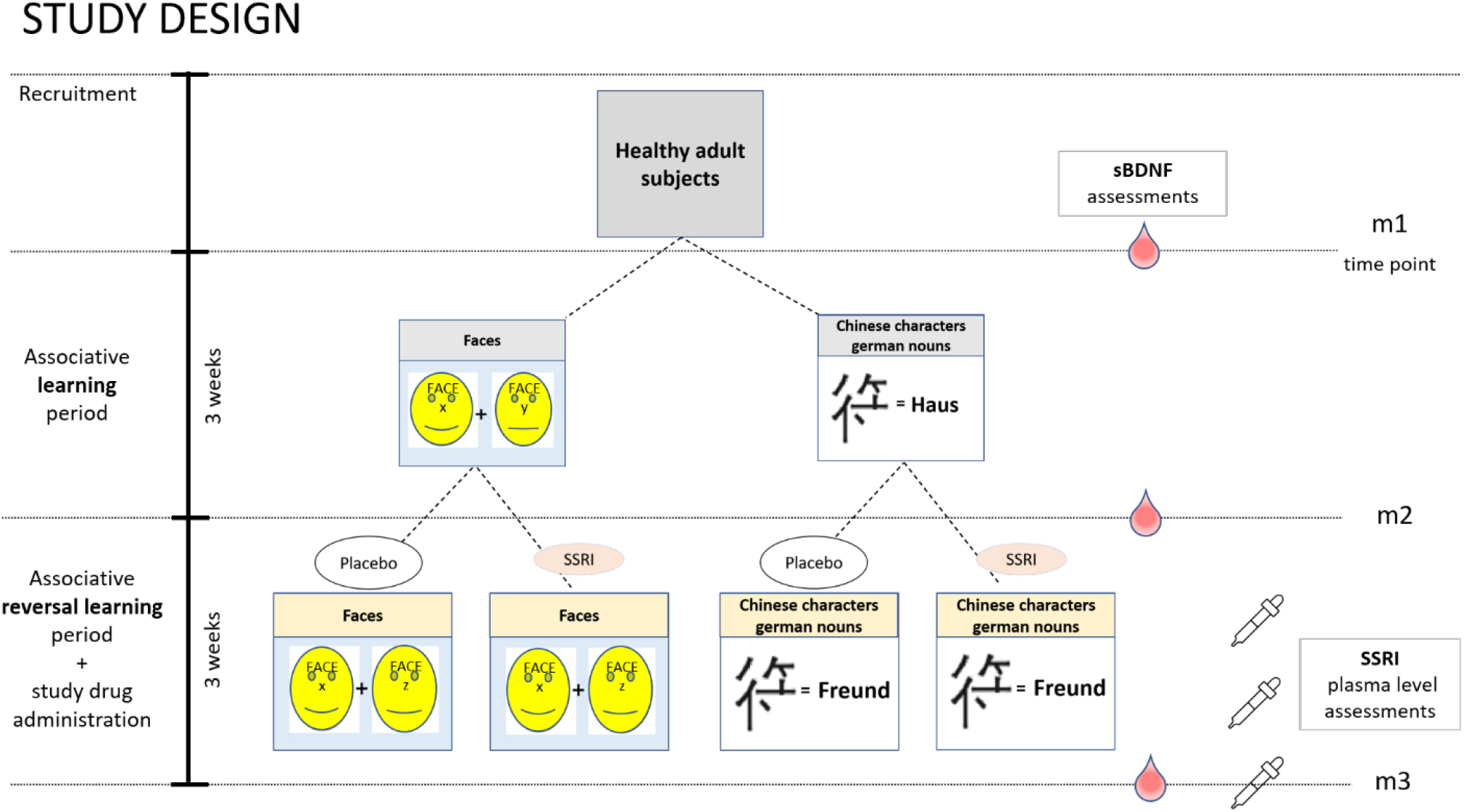
Study design. After inclusion, subjects were randomly assigned to one of two learning paradigm groups, either association of face pairs (emotional content group) or association of Chinese characters to unrelated German nouns (semantic content group). After a period of three weeks with daily learning, each group was again randomly split into a placebo group and an SSRI group (10mg escitalopram orally per day). Subsequently, all four groups underwent another three weeks of associative reversal learning, where either face pairs or characters/nous had changed. Serum levels of BDNF were assessed at three time points: before (m1) and after the three-week associative learning period (m2), as well as after the three-week reversal learning period (m3). Additionally, citalopram plasma levels were assessed 7, 12 and 21 days after beginning of administration; at m3 SSRI level and BDNF levels were assessed simultaneously.

### Subjects

Data from 37 healthy, non-related adult subjects were included into the analysis (mean age ± SD: 25.92 ± 4.97 years, 20 females, 17 males). General health was assessed by means of medical history, physical examination, electrocardiography and blood sampling. To screen for psychiatric health, a structured clinical interview for DSM-IV (SCID) was conducted. Moreover, inclusion criteria were willingness and competence to participate in the study, age between 18 to 55 years and non-smoking status. Exclusion criteria comprised medical, psychiatric, or neurological illnesses, a lifetime use of SSRIs, first degree relatives with a history of psychiatric illness, colour blindness, non-Caucasian ancestry (self-reported) and knowledge of Kanji or Hanzi (Japanese or Chinese characters). All subjects gave written informed consent and received reimbursement for their participation.

### Associative learning paradigms

Associative learning comprised memorising either two faces to one another or Chinese characters to unrelated German nouns. Faces for the emotional content learning group were taken from the “10k Adult Faces Database”[18]. Chinese characters for the semantic content learning group were randomly selected and had no meaning or connection to the associated words. Per day, 52 images pseudo-randomly selected out of 200 pairs were presented to the subjects at an online platform that was developed in house. Each image was presented for 5 seconds. Each session was followed by a retrieval. From the subset of all previously learnt associations one half of the random pairs was shown, while the correct association had to be chosen from four possible answers with unlimited time. Subjects did not receive any feedback on their learning performance.

### Assessment of serum BDNF levels

Serum BDNF levels were measured from peripheral venous blood samples. Samples were collected in serum vacutainer tubes (Becton Dickinson). Subsequently, tubes were centrifuged at 1500 x g for 15 min. The liquid portions were pipetted and stored at -80°C before analysis of sBDNF levels. Assessment of sBDNF levels was performed using an ELISA (enzyme-linked immunosorbent assay) kit (Biosensis® Mature BDNF Rapid™ ELISA Kit: Human, Mouse, Rat; Thebarton, SA, Australia). As described in the manufacturer’s protocol, serum samples were appropriately diluted (1:100) and BDNF was detected on pre-coated mouse monoclonal anti-mature BDNF 96-well plates. Within five minutes after addition of stop solution, absorbance was measured in a microplate reader set at 450 nm and a correction wavelength set to 690 nm to determine BDNF concentrations according to the standard curve. The assays were performed in duplicate and the mean of both was calculated. For analysis of intrinsic assay quality, intra- and inter-assay coefficients of variation (CV) were assessed.

### Study drug administration and assessment of SSRI plasma levels

For 21 days, subjects received either a placebo or 10mg escitalopram (Cipralex® Lundbeck A/S, provided by the Pharmaceutical Department of the Medical University of Vienna) per day. With escitalopram we chose a study drug that has a specific effect on the serotonergic system by selectively binding to the transporter. Escitalopram possesses an adequate tolerability and dose-dependent mild and temporary adverse events[19]. Previous studies had already shown its tolerability in healthy volunteers when administered longitudinally[20]. Plasma levels for escitalopram can be detected from citalopram plasma levels since the latter contains both, S- and R-citalopram. Venous blood samples were drawn 7, 14 and 21 days after administration onset. Plasma levels were measured at the Clinical Department of Laboratory Medicine of the Medical University of Vienna using liquid chromatography–tandem mass spectrometry (LC-MS/MS). For citalopram the therapeutic reference range is 50-110 ng/ml, for escitalopram (S-citalopram) the reference range is 15-80 ng/ml[21].

### Statistics

To assess the effects of learning and SSRI administration on sBDNF levels, linear mixed effects models (LMEs) were used. The learning (m1 and m2) and relearning phases (m2 and m3) were analysed separately. For the learning model, “group” (characters / faces) and “measurement” (m1 / m2) were included as factors. The relearning model contained an additional “substance” factor (placebo / escitalopram). Interactions between these factors were assessed and dropped if not significant. “Age” (mean-centred) and “sex” were always added as covariates. A second relearning model additionally corrected for the final citalopram plasma levels (mean-centred) was also constructed. All models contained a random intercept per subject. A random “measurement” slope per subject was added only if it increased the model quality (as indicated by the Akaike information criterion). All factors were reference coded as follows, for the learning model m1, and for the reversal learning m2 was used as reference measurement. For the other factors, Chinese character and placebo groups were used as references. Finally, to test for an association between sBDNF values and SSRI plasma levels, partial Pearson correlation was performed, controlled for “age”, “sex” and “group”. Additionally, partial Spearman was performed, to account for outlying values. For all analyses, alpha threshold of significance was set at p<0.05. Calculations were performed using MATLAB 2018b.

## 3. Results

### Subgroups

Data from 37 subjects were included in the final analysis. For group sizes see table 1.

**Table 1.**
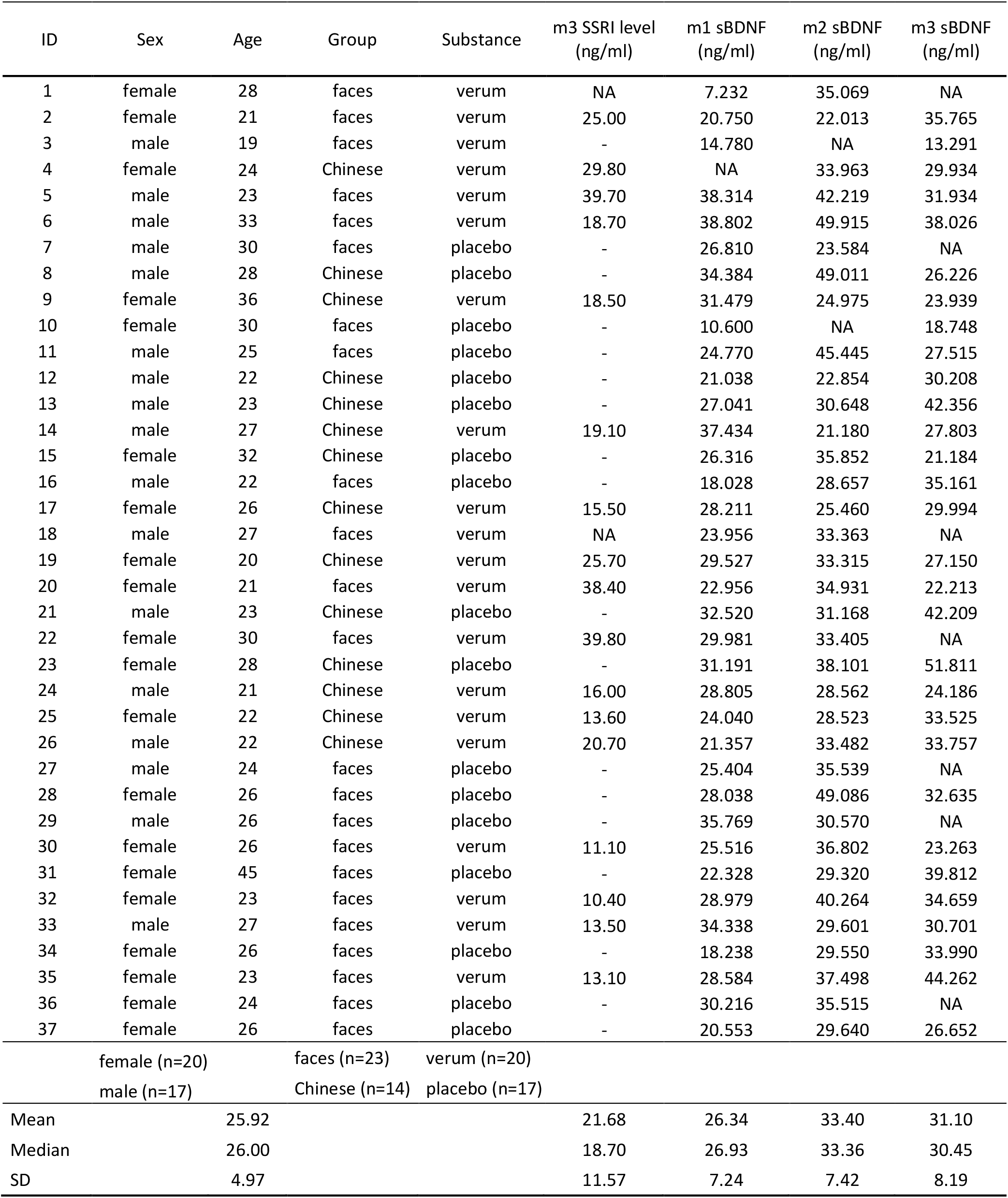
Subjects, group assignments and acquired data. The table lists the full data for the 37 subjects included in the analysis. Values for mean plasma level, median and SD only refer to the verum group. Due to blinding, SSRI levels were also assessed in the placebo group. Values below the detectability threshold were indicated as “–”. Two subjects (ID 1 and 18) dropped out before m3. NA: not available, SD: standard deviation, sBDNF: serum brain-derived neutrophic factor, SSRI: selective serotonin reuptake inhibitor

**Table 2.** LME results for interaction effects of learning period (m1 / m2) and groups (faces / Chinese) on sBDNF levels. DF: degrees of freedom, CI: confidence interval, SE: standard error, m2: measurement 2, faces: emotional content learning group. Intercept refers to male, m1 and character.

| Name | Estimate | SE | t-value | DF | p-value | 95% CI |
| --- | --- | --- | --- | --- | --- | --- |
| Intercept | 30.14 | 2.09 | 14.45 | 65 | 0.00 | [25.97, 34.30] |
| m2 | 2.53 | 2.33 | 1.09 | 65 | 0.28 | [-2.12, 7.18] |
| female | -2.68 | 1.99 | -1.35 | 65 | 0.18 | [-6.64, 1.29] |
| faces | -3.67 | 2.33 | -1.58 | 65 | 0.12 | [-8.32, 0.97] |
| age | 0.16 | 0.20 | 0.80 | 65 | 0.43 | [-0.24, 0.56] |
| m2 ~ faces | 6.88 | 2.97 | 2.32 | 65 | 0.02 | [0.96, 12.81] |

**Table 3.**
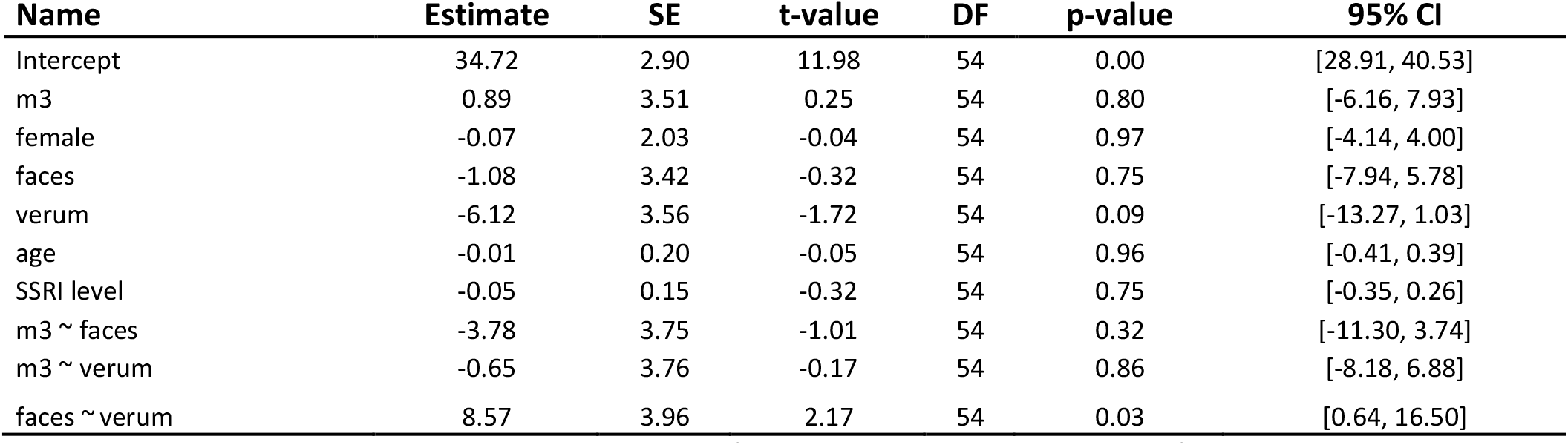
LME results for interaction effects of substance (verum / placebo) and reversal learning (faces / Chinese) on sBDNF levels. DF: degrees of freedom, CI: confidence interval, SE: standard error, m2: measurement 2, faces: emotional content learning group. Intercept refers to male, m2, character and placebo.

### sBDNF levels at baseline and during the study

At the first measurement (m1), mean sBDNF levels for all subjects were 26.34 ± 7.24 ng/ml (mean± SD), followed by 33.40 ± 7.42 ng/ml at the second measurement (m2), and 31.10 ± 8.19 ng/ml at the third measurement (m3), see table 1.

### Effects of learning period and groups on sBDNF levels

For the learning phase, a significant interaction between “measurement” and “group” was found (p = 0.02) indicating a higher sBDNF level in the “faces” group at m2.

### Effects of substance and reversal learning paradigms on sBDNF levels

No three-way interaction between “measurement”, “group” and “substance” was detected, independent of a correction for the final citalopram levels. A “group-by-substance” interaction was found with (p = 0.03) and without (p = 0.03) the SSRI level as covariate. The covariate for the final citalopram levels was not significant, which is also reflected in the similar p-values.

### Correlation between sBDNF levels and SSRI plasma level after three weeks of SSRI administration

Partial Pearson correlation (r=-0.30, p=0.05) revealed a negative association between SSRI and sBDNF level after three weeks of escitalopram administration and relearning. Partial Spearman correlation also showed a negative, albeit less pronounced, dependency (r=-0.22, p=0.15) (Figure 2).

**Figure 2.**
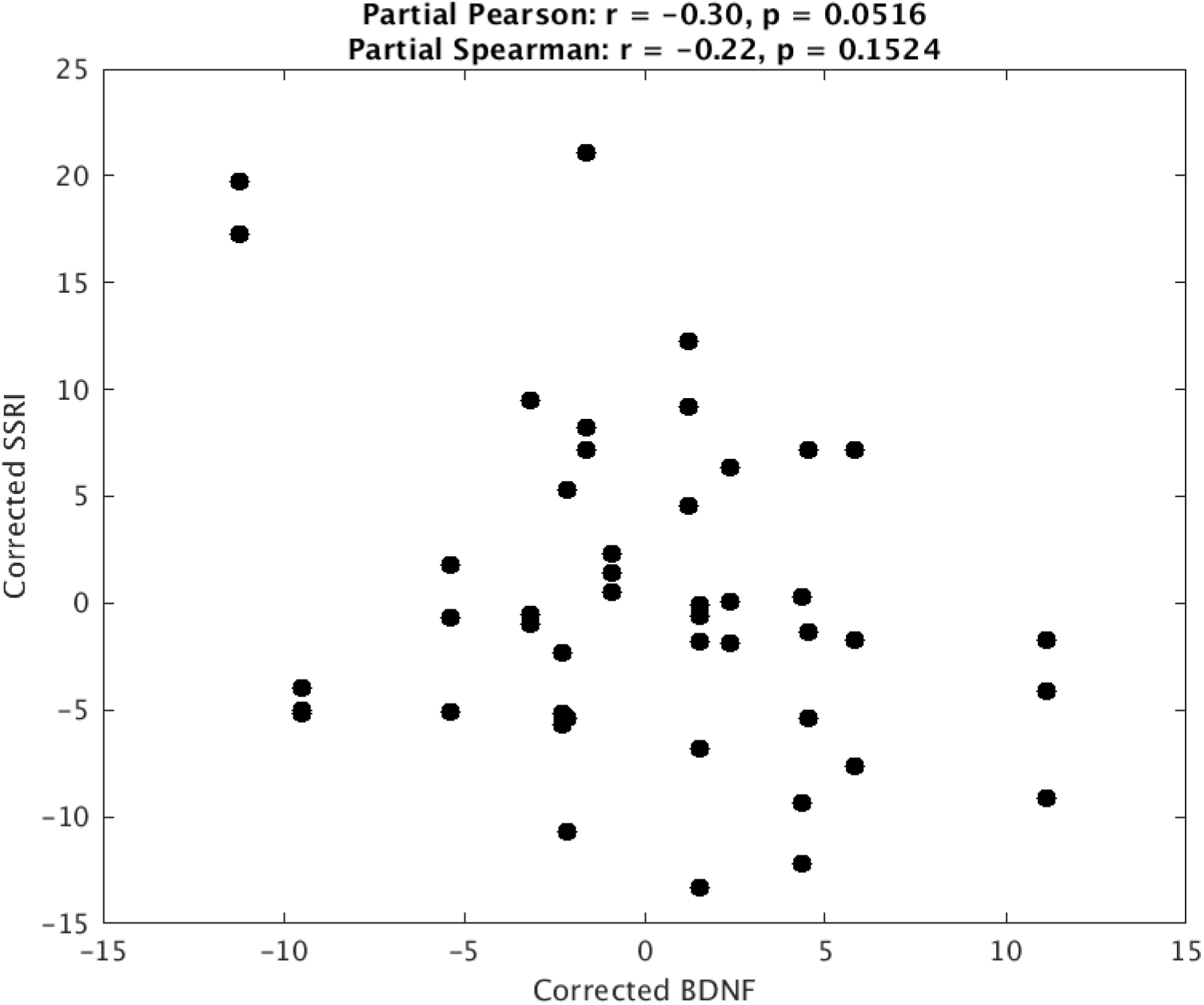
Correlation of sBDNF and SSRI plasma levels at the end of the relearning phase. Partial Pearson correlation of sBDNF and SSRI level, controlled for age, sex and group, indicates a significant negative correlation after a three-week period of daily escitalopram administration (10 mg) and the concomitant associative reversal learning period. Partial Spearman’s rank correlation does not reach significance. sBDNF – serum brain-derived neutrotrophic factor, SSRI-selective serotonin reuptake inhibitor

## 4. Discussion

In this randomised, double blinded longitudinal intervention study we aimed to investigate the effect of associative learning and reversal learning in combination with three-week daily escitalopram intake on serum BDNF levels in healthy human subjects. Learning face pairs as compared to associations between Chinese characters and unrelated German nouns lead to a stronger increase in sBDNF.

Learning tasks as those employed here were shown to activate predominantly hippocampal and parahippocampal regions[22, 23]. To increase activation of limbic regions, including the hippocampus, emotional content was added to our learning tasks[24, 25]. In mice, BDNF seems to be necessary for specific learning tasks that implicate the hippocampus[9]. Studies on BDNF and learning in healthy subjects, however, are scarce. The association of peripheral BDNF with cognitive functioning is often investigated in combination with physical activity[26], where BDNF serves as a marker of plasticity. Moreover, while BDNF levels seem to correlate with better memory performance, no difference was found through repeated cognitive training. As an explanation, it was suggested that higher cognitive performance might require higher BDNF levels[27]. According to the *synaptic tag* hypothesis[28], BDNF might represent a plasticity-related protein that is already induced by a weak behavioural training task. This could be represented by the increase in BDNF levels we see in our model in the initial associative learning phase. Moreover, the effect of our emotional content learning paradigm on sBDNF levels could resemble the requirement for BDNF in the hippocampus during the period of emotional learning. BDNF was shown to be influenced by (treatment) interventions such as electroconvulsive therapy[29, 30] and SSRI administration[10]. Treatment with fluoxetine leads to elevated levels of BNDF mRNA in the mouse hippocampus, however, only in combination with fear extinction training[31]. Since habituation also represents a learning phenomenon[32], it can be conceived that our reversal learning task might have been accompanied with too little novelty to induce further changes of peripheral BDNF levels, despite the potentially modulating effect of chronic SSRI administration[12].

The correlation of peripheral BDNF and SSRI levels continues to be a matter of debate. In our study, we observed a trend towards a negative correlation of sBDNF and plasma citalopram levels at the end of the three-week drug administration. Given the magnitude of the effect and the non-significance of the correlations, this result can only be an indication of a possible relationship. Meta-analytical findings point towards an increase in peripheral BDNF levels through antidepressant treatment in patients with depression[10]. Also, BDNF levels were shown to be lower in therapy-naïve patients than in patients with depression treated with antidepressants, or in healthy control[33]. Accordingly, post-mortem studies find higher cortical BDNF expression in SSRI-treated than in untreated depressed patients[34]. Moreover, an increase of BDNF through chronic SSRI administration in the visual cortex and hippocampus of the rat was shown[35]. One proposed mechanism for the effect of SSRIs is an upregulation of CREB mRNA in the hippocampus and subsequently an increased expression of BDNF and TrkB[36].

Knorr et al. have been, until now, the only group to investigate the effect of chronic escitalopram administration on whole blood BDNF levels in healthy subjects[20]. They reported a significant negative correlation between citalopram and whole blood BDNF levels. In line with their results, escitalopram was shown to lead to a decrease of BDNF protein in the rat hippocampus and frontal cortex, while not affecting BDNF mRNA expression in these regions[37]. Further, a decrease of BDNF mRNA in the dental gyrus of rats was demonstrated, which was induced separately by various serotonergic agents including paroxetine, tranylcypromine, and *p*-chlorophenylalanine[38]. Our results seem to mirror these findings, reflecting those central processes in the periphery. Most importantly, as one of only a few studies, our data allows for the direct correlation of drug levels (in this case citalopram) and peripheral BDNF levels, instead of investigating BDNF levels over a treatment period alone. A molecular mechanism for the negative correlation between citalopram plasma levels and BDNF serum levels, however, is missing. The reduction of peripheral BDNF levels with increasing escitalopram levels might be an indication of increased binding of BDNF with a greater TrkB receptor accessibility enabled through the SSRI[12].

Five subjects had a citalopram level beneath the therapeutic reference range. Notwithstanding, a plasma concentration of 2.5 ng/ml citalopram was shown to result in a serotonin transporter occupancy in the midbrain of 50%[39]. In a PET study, our group has shown strong correlations of serotonin transporter occupancy and escitalopram plasma levels after a drug administration of three weeks[40]. Consequently, effects of chronic drug intake on brain metabolism and cognitive functions could be expected according to our citalopram plasma concentrations.

## 5. Limitations

To differentiate between effects of learning and no intervention in general, it would have been beneficial to include an additional non-learning control group. Also, numerous factors affecting sBDNF levels were not accounted for in this study, e.g. physical activity or seasonal variations[41, 42]. Here, a systematic survey of the most important influencing factors would have been beneficial. It should also be mentioned that some subjects exhibited a citalopram plasma level below the therapeutic range. Although medication adherence was controlled using therapeutic drug monitoring[43], it is not possible to definitively evaluate whether this variation is solely attributable to study drug adherence or also to rapid/slow metabolisation. Moreover, the dosage, length of drug administration[10] as well as learning duration[44] might be reconsidered in future studies.

## 6. Conclusion

This study assessed the effects of associative learning and SSRI administration (escitalopram) on serum levels of BDNF, a neurotrophic factor involved in neuroplasticity and prominent target for recent psychiatric research. Our results indicate a modulating influence of emotional compared to non-emotional learning content on BDNF. Furthermore, indications of a negative correlation between SSRI plasma levels and sBDNF were found. While the definite mechanism responsible for this relationship is still missing, the findings present an additional piece of information in the current debate on the dynamic interplay of SSRIs, learning and peripheral BDNF.

## Acknowledgements

We thank T. Stimpfl for SSRI plasma level analyses. Also, we wish to thank the team members as well as the medical students of the Neuroimaging Lab (NIL), headed by Prof. Lanzenberger, the corresponding author.

## Declarations

### (iii) Availability of data and material

The full data can be made available upon request to the corresponding author.

### (iv) Ethics approval

This study is part of a larger project that has been approved by the ethics committee of the Medical University of Vienna (EK Nr.: 1739/2016) and performed in accordance with the Declaration of Helsinki (1964). The project is registered at clinicaltrials.gov with the identifier NCT02753738.

### (v) Consent

All subjects provided written informed consent before inclusion into the study.

### (vi) Author contributions

R. L., R.S. and T. V. designed the main study. M. K., J. U., G.M. G., T. V. established the study concept and data analysis. M. K. performed data analysis. J. U. and G. M. G. wrote the first draft of the document. V. R. and D. P. provided support in recruitment and data collection. N.V. and A. E. performed BDNF analysis. All authors contributed to and have approved the final manuscript.

#### Abbreviations

BDNF: brain derived neurotrophic factor
CNS: central nervous system
CV: coefficient of variation
CYP: cytochrome p 450 polymorphism
CREB: cAMP-response element binding protein
ECG: electrocardiograph
ELISA: enzyme-linked immunosorbent assay
ERK: extracellular signal-regulated kinases
LC-MS/MS: liquid chromatography–tandem mass spectrometry (LC-MS/MS)
LME: Linear mixed effects
LTP: long-term potentiation
MAPK: mitogen-activated proteinkinase
MAO: monoaminooxidase
mRNA: messenger ribonucleic acid
sBDNF: serum BDNF
SCID: structured clinical interview from
DSM-IV SSRI: serotonin reuptake inhibitors
TrkB: tropomyosin-related kinase B

